# Unravelling the role of *IRX4* variants in non-syndromic and Down syndrome associated congenital heart disease

**DOI:** 10.64898/2026.09.07.749900

**Authors:** Jyoti Maddhesiya, Kriti Jain, Dharmendra Jain, Ashok Kumar, Amit Rai, Bhagyalaxmi Mohapatra

## Abstract

*IRX4* is a TALE- homeodomain transcription factor which is essential for cardiac development. In murine models, *Irx4* deficiency leads to impaired ventricular function and results in cardiomyopathy. To elucidate the role of *IRX4* in human congenital heart disease (CHD), Sanger sequencing of the *IRX4* gene was performed in 205 individuals with non-syndromic CHD, 24 Down syndrome (DS) cases with CHD, 27 DS cases without CHD, and 150 healthy control individuals. Two novel (p.Ser24Asn and p.Thr217Iso) and one reported variant (rs2232376) were identified in non-syndromic CHD. Concurrently, rs2232376 was also detected in DS with CHD. The first novel (p.Ser24Asn) and reported (rs2232376) variants lie in the N-terminal region while the second novel (Thr217Iso) variant lies within the TALE homeodomain. *In silico* structural modelling suggested that both the novel variants (p.Ser24Asn and Thr217Iso) induce conformational changes in the IRX4 protein, potentially altering its DNA-binding affinity. A significant reduced expression of IRX4 muteins was noted in Western blotting by both variants (p.Ser24Asn and Thr217Iso). Furthermore, luciferase reporter assays demonstrated decline in the activity of *Nanog* promoter and *HEY2* enhancer in response to both the variants which was further corroborated by decrease mRNA expression in qRT-PCR. Additional downstream targets, including *Nfyc*, *Nppa*, and *Bmp10*, also exhibited anomalous expression due to both the variants (p.Ser24Asn and Thr217Iso). Altogether, the aberrant expression of muteins as well as downstream target genes along with compromised activities of promoters substantiate the pathogenic potential of the identified *IRX4* variants and underscore the critical role of *IRX4* in regulating multiple stages of cardiogenesis.

## 1. Introduction

Congenital heart diseases (CHDs) are the structural malformations of heart that occur during embryonic development, and presented mostly in neonates, accounting nearly 1% in live births (Hoffman, 2002; van der Linde et al., 2011). Considering CHDs are developmental defect, genetic basis of the disease is predominantly predicted, even though influence of epigenetic and environmental factors could not be ruled out. CHDs are genetically complex, manifested with distinct clinical phenotypes, either alongside extracardiac malformations and grouped as syndromic CHDs (sCHDs) or in isolation, which is classified as non-syndromic CHDs (nsCHDs) (Eskedal et al. 2007; Maddhesiya and Mohapatra 2024). Among these, Down syndrome (DS), most frequently occurring syndrome, exhibit co-occurrence of CHD (septal defects) in approximately 40-50% cases along with other extracardiac syndromic features. The cardiac phenotype primarily include atrioventricular septal defect (AVSD), atrial septal defect (ASD), ventricular septal defect (VSD), and tetralogy of Fallot (ToF) (Pradat 1992; Stoll et al. 1998a; Freeman et al. 1998; Maddhesiya and Mohapatra 2024). Pathogenic variants in *CRELD1* gene is strongly associated with CHD-DS, have been identified in several studies (Maslen et al. 2006; Asim et al. 2017; Leiva-Cabrera et al. 2025). Besides *CRELD1*, other genes such as *COL6A1*, *COL6A2*, *FBLN2*, *FRZB*, *GATA5*, *HEY2*, *GATA3*, *RCAN1*, *GUSB*, *KCNH2*, *CEP290*, and *ENG* have also been implicated in CHD (Gelb and Chung 2014; Alharbi et al. 2018; Moran and Robin 2020). Despite extensive research during past 50 years, genetic correlation among nsCHD and DS-CHD is not fully understood.

*IRX4*, also known as Iroquois homeobox protein 4, encoded by *IRX4* gene, well recognised to be involved in embryonic development and cellular differentiation. It is a ventricle-restricted transcription factor and an early marker of ventricular progenitors, maintaining conserved ventricular expression across multiple species (Bao et al. 1999; Bruneau et al. 2000; Wang et al. 2001; Garriock et al. 2001). Interestingly, its expression is significantly diminished in *Nkx2.5^−/−^* and *dHand^−/−^* mouse embryos (Bruneau et al. 2001; Anderson et al. 2018). Loss of *Irx4* leads to abnormal ventricular gene expression and cardiomyopathy in mice (Bruneau et al. 2001). Functionally, *Irx4* activates *VMHC1* and indirectly suppresses *AMHC1*, while Irx4-null mice show elevated cardiac stress markers such as *BNP*, α*-actin*, and β*-MHC* (Kim et al. 2012). *Irx4* knockdown reduces cell proliferation, sphere-forming capacity, and expression of stemness/ differentiation markers including *CD133*, *Aldh1A1*, *Nanog*, *Sox2*, and *Notch1* (Jia et al. 2020). Combined loss of *Irx3* and *Irx4* results in increased heart failure markers, enhanced Bmp10 signaling, and irregular cardiomyocyte proliferation, features consistent with left ventricular non-compaction (LVNC) (Liu 2017). In *Smyd1*-null mice, both *Hand2* and *Irx4* levels are reduced in the developing heart by embryonic day 9.0 (Gordon et al. 2022). Only one study to date reporting a link between non-synonymous variants in *IRX4* and nsCHD (Cheng et al. 2011). Despite its critical roles in cardiac development, the involvement of *IRX4* in CHD remains poorly understood.

In light of this, the current study was designed to evaluate the role of *IRX4* gene, by identifying variants in *IRX4,* in both non-syndromic CHD and CHD associated with DS in the north Indian population, and to assess the functional consequences of these variants. Sanger sequencing of the *IRX4* gene revealed two novel missense variants as well as one previously reported variant in nsCHDs. The known variant was also detected in DS-CHD. *In silico* and *in vitro* analyses depicted that both novel variants likely impair the DNA binding affinity of the IRX4 protein, which could disrupt the coordinated cardiac regulatory network involved in cardiogenesis, thereby contributing to the observed CHD phenotypes.

## 2. Materials and Methods

### 2.1 Recruitment and clinical evaluation of the subjects

This is a case-control study, performed after enrolling 205 non-syndromic CHD (nsCHD), 24 DS with CHD, 27 DS without CHD probands and 150 healthy age-matched controls from the same geographical location. All the subjects were recruited from Department of Cardiology and the Department of Paediatric Medicine, SS Hospital, Banaras Hindu University (BHU), Varanasi, India after a detailed written informed consent before sample collection. The enrolled individuals were thoroughly investigated clinically by 2D-colour doppler echocardiography, chest radiography, and ECG while DS patients were confirmed by Karyotyping along with echocardiography. The study designed was approved by the ‘Institutional Human Ethical Committee’ of the University.

### 2.2. Genetic screening, identification, and analysis of variants

Genomic DNA was extracted from the collected peripheral blood samples, by standard ethanol precipitation protocol. Both the protein coding regions as well as the flanking exon-intron boundaries of human *IRX4* gene (NM_001278635.2) were amplified by polymerase chain reaction (PCR) with the non-syndromic genomic DNA and reaction mixture (10X PCR buffer (2 μl), 50 mM MgCl_2_ (0.8 μl), 10mM dNTP mix (0.5 μl), 10 μM forward and reverse primers (0.2 μl each) and Taq DNA polymerase (1U) with sterile nuclease-free distilled water for volume adjustment). The PCR products were enzymatically purified by Exonuclease I (USB Products, Affymetrix, Inc., USA) and recombinant Shrimp Alkaline Phosphatase (USB Products, Affymetrix, Inc., USA). Sequencing of purified amplicons were carried out by Sanger sequencing method using Big Dye® Terminator v3.1 Cycle Sequencing Kit (Applied Biosystems, Inc. USA) on genetic analyser by Applied Biosystems (ABI PRISM 3500, USA). Using Finch TV software (http://www.geospiza.com/ftvdlinfo. html, Geospiza), the chromatogram of sequence was analysed. Each identified variant was confirmed by re-sequencing of the variation carrying DNA samples with alternate primer as well as with independently amplified PCR amplicon from the respective proband. The inheritance pattern was checked by the parental genotyping. Various databases namely ClinVar (http://www.ncbi.nlm. nih.gov/clinvar/), gnomAD (https://gnomad.broadinstitute.org/), and 1000G (https://www.internationalgenome.org/) were searched for novelty of variants.

### 2.3 Computational analysis

To predict the pathogenicity of identified missense variants on the structure, stability, accessibility, and function of proteins various online *in silico* tools were queried. To proceed with the described analyses, *IRX4* reference genomic DNA and mRNA sequence from Gen-Bank (https://www.ncbi.nlm.nih.gov/genbank/) and protein sequence from Protein database (https://www.ncbi.nlm.nih.gov/protein/) were retrieved.

#### 2.3.1 Phylogenetic conservation analysis

To check the evolutionary conservation of mutated amino acid across various orthologues species of vertebrate including *H.sapiens, P.troglodytes, M.mulatta, C.lupus, B.taurus, M.musculus, R.norvegicus, G.gallus, D.rerio,* multiple sequence alignment was performed using the ‘Homologene’ feature of NCBI (http://www.ncbi.nlm.nih.gov/homologene).

#### 2.3.2 Potential pathogenicity prediction

The pathogenic potential of the identified variants was predicted by using an integrated database Varcard2 which incorporate more than 10 bioinformatic tools such as Polyphen2_HDIV, MutationTaster, PROVEAN, VEST3, M_CAP, CADD, DANN, GenoCanyon, ClinPred, ReVe, etc. The prediction of these tools based on different parameters and algorithms such the conservation of nucleotides and conservation of amino acid residues as nature of R-group.

#### 2.3.3 Prediction of RNA Secondary Structures

Using the RNA Structure Web Server (version 6.0.1) (http://rna.tbi.univie.ac.at/cgi-bin/RNAWebSuite/RNAfold.cgi), the secondary structures and stability of RNA was predicted (Bellaousov et al. 2018). The server is based on the algorithms of thermodynamic principles to forecast RNA secondary structures. The folding patterns and potential base pairing interactions within the RNA molecules can be speculated.

#### 2.3.4 Mutational effect on RNA features

To further explore the structural alterations induced by each missense mutation, MutaRNA tool was used (Miladi et al. 2020). The analysis involved the intra-molecular base pairing potential, base pairing probabilities, and RNA accessibility (single-strandedness) of the mutant (MUT) mRNA when compared to wild-type (WT). By incorporating remuRNA (Salari et al. 2013) and RNAsnp, the structural alterations induced by mutations are better understood.

#### 2.3.5 Prediction of physicochemical properties

Missense variants also affect the physicochemical properties of proteins viz, relative mutability, recognition factors, total beta strand and hydrophobicity etc. As the R-side chains of each amino acids regulate these properties and alteration induced by missense variations, ultimately change the above-mentioned properties of mutant proteins (muteins). Using a web-based server Protscale (https://web.expasy.org/protscale/), a relative study was performed to check the effect of mutations on these physicochemical properties.

#### 2.3.6 2-D and 3-D conformational changes in muteins

To predict the alterations in the secondary structures of muteins, an online server Psipred (http://bioinf.cs.ucl.ac.uk/psipred/) was searched. Additionally, the tertiary structures were predicted using Alphafold2 googlecolab (https://colab.research.google.com/github/sokrypton/ColabFold/blob/main/AlphaFold2.ipynb), the conformational changes in the tertiary structures of muteins induced by the missense variants were also predicted. To visualize the structural changes in the modelled structures, PyMOL Molecular Graphics System, Version 3.1 (Schrödinger, LLC.) was used followed by RMSD value calculations.

### 2.4 Functional characterization of variants by in vitro analysis

#### 2.4.1 Cloning followed by site-directed mutagenesis

The expression clone of *Homo sapiens IRX4* with full-length ORF and tagged with mCherry was purchased from “Addgene” (Cat No. #72692, Watertown, MA, USA). The mutant constructs (IRX4_S24N and IRX4_T217I) were prepared for the novel identified variants (p.Ser24Asn and p.Thr217Iso) using ‘site directed mutagenesis kit’ (Agilent Technologies Inc.), and with the help of a complementary pair of designed mega primers. Transformation of the mutant constructs was performed into DH5α competent cells followed by plasmid isolation (Qiagen midi kit, Germany). To validate the MUT constructs, Sanger sequencing was performed.

#### 2.4.2 Cell culture and transfection

In order to conduct *in vitro* functional analysis, mouse pluripotent embryonic carcinoma cell line P19 (kind gift from Dr. Ramkumar Sambasivan, InStem, India) and cadiomyoblast H9c2 cells (purchased from NCCS, Pune, India) were used. P19 cell line was cultured in α-MEM (Gibco, Life technologies Corp.) with 10% fetal bovine serum supplements (Gibco, Life technologies Corp.) and 100 units/mL penicillin and 100 units/mL streptomycin while H9c2 (due to cardiomyoblast in origin and with better morphology, mainly used for immunostaining experiment) incubated in Dulbecco’s modified Eagle’s medium (DMEM) with 10% serum supplements at 37°C with 5% CO2 in a humified chamber. For each experiment, transfection was carried out at 30-40% confluency with FuGENE 6 transfection reagent (Promega Corp., IN, USA) after 24 hrs of cells seeding.

#### 2.4.3 Western blotting

The effects of variants on the expression of IRX4 proteins was estimated by Western blotting. P19 cells were seeded in 6 well-plates followed by transfection with IRX4_WT and MUT constructs (1 µg) after 24 hrs. Post 48 hrs of transfection, whole cell lysate was prepared in RIPA Buffer (as per Dixit et al, 2021) and isolated protein was quantified by Bradford Assay. 25 ug of protein were diluted in Laemmeli’s loading buffer followed by denaturation for 5 mins at 95°C and then lysates were resolved on 8% SDS polyacrylamide gel. Proteins were transferred to PVDF membrane (Bio-Rad Laboratories Inc, CA, USA) and blocked in 5% dry skimmed milk in 1X TBST (25 mM 1 M Tris-Cl pH = 7.5, 0.15 M NaCl with 0.1% Tween 20) for 2 hrs at room temperature (RT) followed by incubation with IRX4-specific primary antibody (Invitrogen), overnight at 4°C. Afterward, the membrane was washed five times with 1X TBST (5 mins per wash), followed by incubation with HRP-conjugated goat anti-rabbit IgG antibody (Genei, Merck Specialties Pvt. Ltd., Germany) for 2 hrs at RT. Subsequently, the membrane was washed 10 times with 1X TBST (5 mins per wash) and visualized using an enhanced chemiluminescence (ECL) detection kit (Amersham, GE Healthcare, USA).

#### 2.4.4 Immunocytochemistry

To check the cellular localization and expression of IRX4 MUT vs WT proteins, immunocytochemistry was performed in H9c2 cell lines. The cells were seeded in 6-well culture plate with each well containing glass cover slips and grown in DMEM supplemented with 10% FBS and incubated in 37°C at 5% CO2. Post 24 hrs, IRX4_WT and MUTs constructs (1 µg) were transfected. Cells were washed with chilled 1X PBS after 48 hrs of transfection, followed by fixation with 4% paraformaldehyde (PFA) immediately for 15 mins. For permeabilization, the fixed cells were treated with 0.5% Triton-X followed by 30 mins of incubation. Cells were washed thrice with chilled 1X PBS and stained with DAPI (Sigma-Aldrich, USA) and Phalloidin (Sigma-Aldrich, USA). Finally, mounted in mounting media and confocal scanning was performed using Zeiss LSM 780 Meta Laser Scanning Super Resolution Microscope System, Germany. Images were analysed by ImageJ, and assembled by Adobe Photoshop software.

#### 2.4.5 Transactivation assay

The transcriptional activity of direct downstream target gene promoters viz, *HEY2-luc* and *Nanog-luc* were checked by dual luciferase reporter gene assay. P19 cells were seeded in 24-well culture plates and transfection was performed after 24 hrs of cell seeding with IRX4_WT and MUT constructs (125 ng) along with each reporter plasmids (125 ng) and Renilla luciferase vector, pRL-TK vector (25 ng) (Promega, USA) as an internal control. Cells were washed with chilled 1X PBS followed by lysates preparation in 1X passive lysis buffer (PLB) after 48 hrs of transfection. Luciferase activity was measured by Dual-Luciferase Reporter Assay System (Promega Corp, WI, USA) using Synergy/ HTX multi-mode reader (BioTek Instruments, Inc., USA). Three independent experiments were performed in triplicate for each sample (WT and MUTs) for both the reporter plasmids and the data were pooled and plotted as mean fold change with standard error of mean. The significance of the data was tested by one-way ANOVA followed by Dunnett’s post-hoc test. Luciferase reporter plasmids used were kind gifts (***HEY2-luc*** from Yusuke Watanabe National Cerebral and Cardiovascular Center Research Institute, Suita, Osaka, Japan and ***Nanog-luc*** from Ariel Waisman, Institute of Neurosciences (INEU), CONICET, Buenos Aires, Argentina).

#### 2.4.6 Real-time PCR (qPCR)

To conduct RT-PCR assay, total RNA was isolated from P19 cells transfected with either IRX4 WT and MUTs constructs, using TRI Reagent (Merck, Germany). The integrity of RNA was checked on 1% agarose gel and further quantified by using NanoDrop (ThermoFisher Scientific, USA). 2 µg of total RNA was treated by DNase I (ThermoFisher Scientific, USA). DNase I-digested RNA was used to prepare cDNA library using Revertaid First Strand cDNA synthesis kit (ThermoFisher Scientific, USA) and Random Hexamers. The assay was performed to evaluate the expression of downstream cardiac enrich genes *Nanog*, *Hey2*, *Nfyc*, *Nppa* and *Bmp10* using real-time PCR machine (QuantStudio 5, Applied Biosystems, CA, USA) and SYBR^Tm^ green (Sigma-Aldrich, USA). Three sets of experiments were performed in triplicate and the pooled data were plotted as fold change with standard error of mean.

### Statistical Analysis

One-way ANOVA test was performed using ‘Graph pad Instat3’ software (GraphPad Software, La Jolla California USA) for analysing luciferase assay and real time PCR data followed by Dunnett’s post-hoc test. *p value* of <0.05 was considered statistically significant.

## 3. Results

### 3.1 Identification of genetic variants and their genotype-phenotype correlation

The coding regions of *IRX4* was screened in 205 non-syndromic sporadic cases of CHD (nsCHD), 24 DS cases with CHD and 27 DS cases without CHD. Sanger sequencing results revealed two novel missense variants (c.71G>A, p.Ser24Asn and c.650C>T, p.Thr217Iso), one reported SNP [c.355G>A, p.Ala119Thr (rs2232376)], 5 synonymous and 6 intronic variations. The first novel heterozygous missense variants (p.Ser24Asn) was identified in 3 unrelated probands (Table 1), which was confined to the N-terminal domain, while the second variant (p.Thr217Iso) lie in the TALE-homeobox domain (TALE-HD) also in heterozygous condition. Both the variants are novel as these are not addressed in any of the clinical database, therefore submitted to ClinVar and their allotted RCV numbers are RCV003318483.1 and RCV003318484.1 respectively. The reported SNP (p.Ala119Thr) confined to the N-terminal domain. None of these variants were detected in 150 healthy age, ethnic-matched control (300 chromosomes). Among the 3 probands harbouring variant (p.Ser24Asn), one 27 yrs old male was diagnosed with VSD while another female is 3 yrs old affected with double outlet right ventricle (DORV), tricuspid atresia (TA) and ASD phenotype, while one more female is 8 months old with cyanotic CHD. The TALE-HD rare-variant (p.Thr217Iso) was detected in 10 months old female associated with VSD. The parental genotype of the male patient carrying p.Ser24Asn variant, was homozygous WT allele (GG) for both mother and father (Table 1). The parental DNA samples of other two cases were not available. In case of variant p.Thr217Iso, only paternal DNA was available, which showed WT (CC) genotype. Although the SNP (p.Ala119Thr) was recognized in 38 CHD probands with highly variable phenotypes *viz.,* VSD (18), ASD (10), Bicuspid aortic valve BAV (5), VSD with patent ductus arteriosus (PDA) (2), two DS-CHD cases also exhibited this SNP. This SNP was identified in a 1 yr old male diagnosed with ASD with DS while the second DS proband harbouring this SNP is affected with ASD and VSD. However, screening of control samples also detects this SNP in our control cohort (100% control). Since the frequency of this SNP is much higher and also detected in controls, functional assessment of this variant was not performed further. The coding positions of substituted nucleotide and amino acid with the clinical CHD phenotype with minor allele frequency (MAF) are presented in Table 1. The sequence chromatogram representing the above listed missense variants and its position on domain of protein are shown in Figure 1(a) and (b).

**Figure 1.**
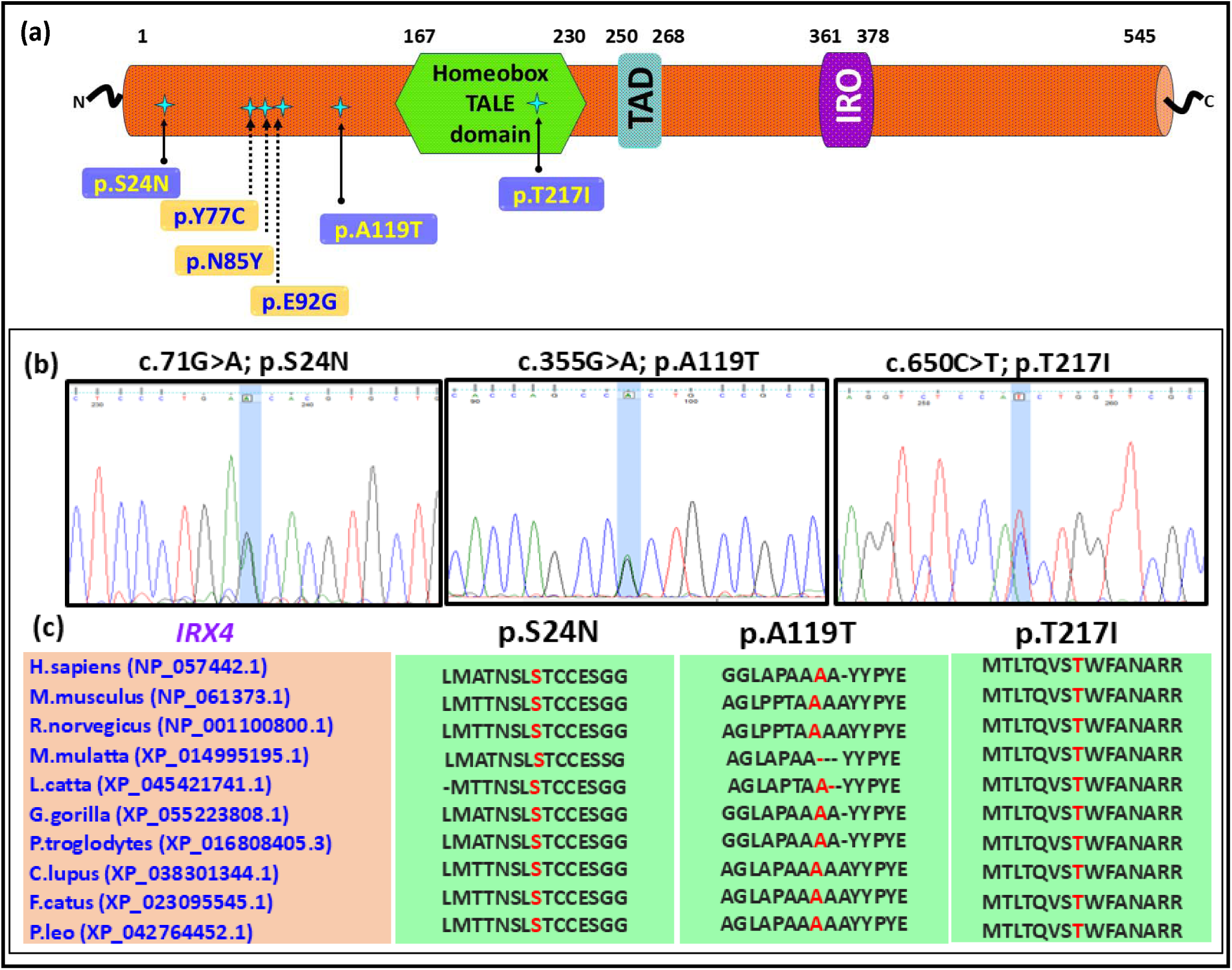
**(a)** Schematic representation of IRX4 which is a 545 amino acid protein with various functional domains viz., homeobox KN domain (187-225aa), Transactivation Domain (235-253aa) and IRO domain (361-378aa). Domain-wise positioning of all the missense identified and previously reported variants (indicated with star) **(b)** the sequence chromatogram peak for the identified variants and **(c)** cross-species alignment of mutated amino acid illuminated in red) across different species of vertebrate for both the variants (p.Ser24Asn and p.Thre217Iso).

**Table 1.**
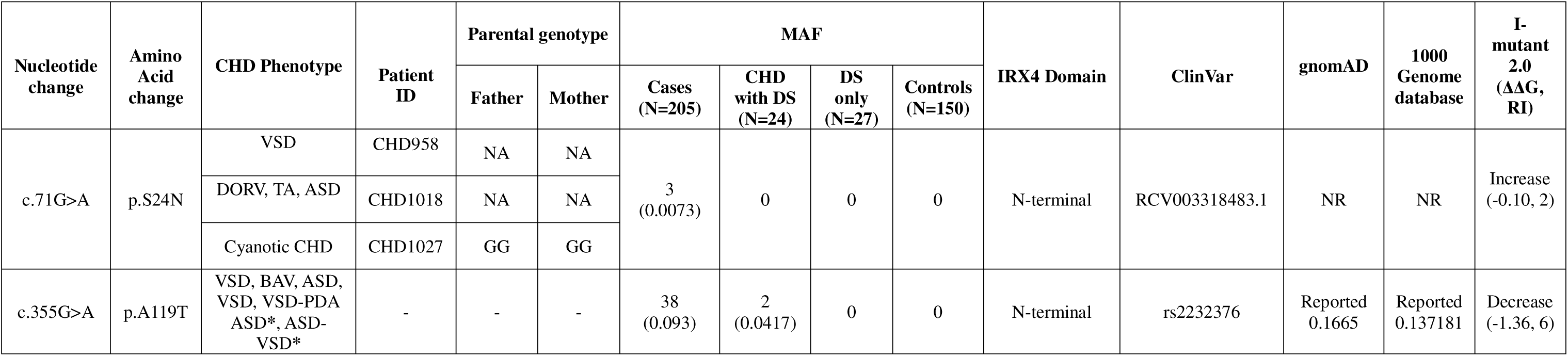
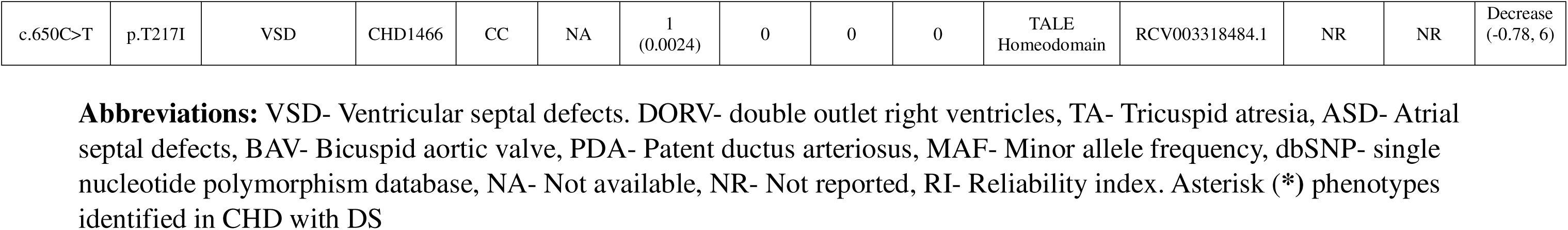
Clinical phenotypes, variants position in domain, ClinVar submission Ids and novelty status.

### 3.2 Analysing phylogenetically conserved IRX4 proteins

In order to check the phylogenetic conservation of the mutated amino acid residues, multiple sequence alignment across various mammalian species such as *Homo sapiens* (NP_057442.1), *Mus musculus* (NP_061373.1), *Rattus norvegicus* (NP_001100800.1), *Macca mulatta* (XP_014995195.1), *Lemur catta* (XP_045421741.1), *Gorilla gorilla* (XP_055223808.1), *Pan troglodytes* (XP_016808405.3), *Canis lupus* (XP_038301344.1), *Felis catus* (XP_023095545.1), *Panthera leo* (XP_042764452.1), revealed 100% evolutionary conservation of the substituted amino acid residues at positions Ser24 and Thre217 are across the above listed species. The evolutionary conserved amino acids with their flanking amino acid sequences have been presented in Figure 1(c).

### 3.3 Prediction of disease-causing potential

The pathogenic potential of both the variants were predicted by Varcad2 server. Almost all the tools (Polyphen2_HDIV, M_CAP, CADD, Mutation Taster, DANN, GenoCanyon and ReVe) speculated that both the variants (p.Ser24Asn and p.Thr217Iso) are damaging and disease-causing. The predictions and the algorithmic scores of both the variants are shown in Table 2.

**Table 2.** Prediction of pathogenic potential of *IRX4* missense variants using different *in-silico* tools.

| Variants | Polyphen2_HDIV |  | M_CAP |  | CADD |  | Mutation Taster |  | DANN |  | GenoCanyon |  | ReVe |  |
| --- | --- | --- | --- | --- | --- | --- | --- | --- | --- | --- | --- | --- | --- | --- |
|  | Prediction | Score | Prediction | Score | Prediction | Score | Prediction | Score | Prediction | Score | Prediction | Score | Prediction | Score |
| c.71G>A | Possibly_damaging | 0.816 | Damaging | 0.214 | Damaging | 22.2 | Disease_causing | 0.913 | Damaging | 0.992 | Damaging | 1 | pathogenic | 0.87427151 |
| c.650C>T | Probably_damaging | 0.999 | Damaging | 0.189 | Damaging | 27.3 | Disease_causing | 1 | Damaging | 0.999 | Damaging | 1 | pathogenic | 0.99766278 |

**Abbreviations:** CADD- Combined annotation dependent depletion, DANN- Deleterious annotation using neural networks

### 3.4 Prediction of secondary structures of RNA

The alterations in the secondary structures of mRNA portrayed a significant changes in the structures induced by both the missense variants (p.Ser24Asn and p.Thr217Iso). The structural modifications are validated by the change in free energy i.e., δδG (-0.8kJ/mol and - 0.9kJ/mol due to p.Ser24Asn and p.Thr217Iso variants respectively). It is speculated that these structural changes affect the RNA stability and folding (Figure 2i(a1-a4).

**Figure 2.**
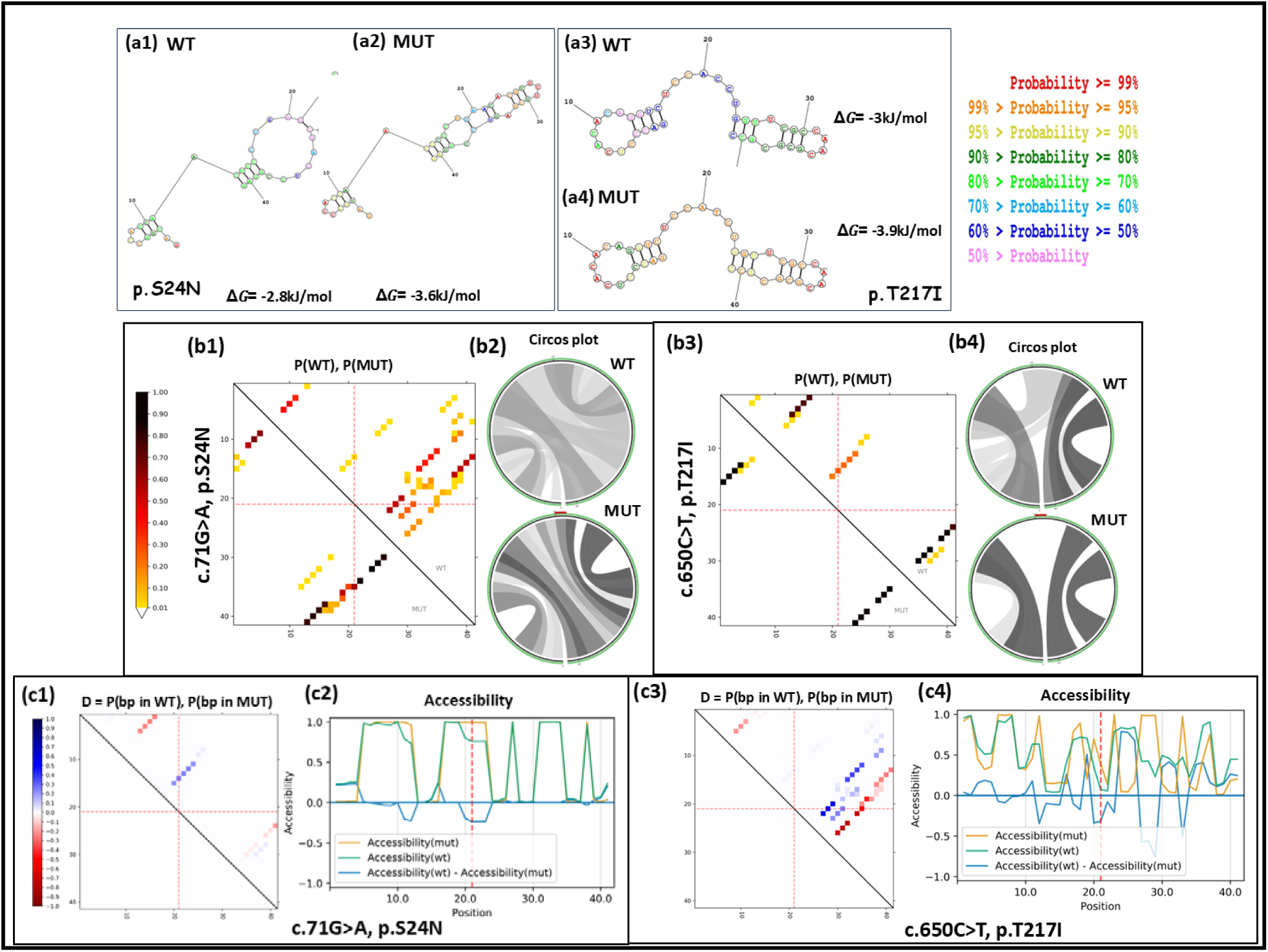
**i(a1-a4)** Predicted secondary structures of WT and MUT RNA for both the IRX4 missense variants c.71G>A (p.Ser24Asn) and c.650C>T (p.Thr217Iso). The mutated nucleotide is at 21^st^ position in each structure. Different base pairing probabilities are denoted by various colors. **ii(b1-b4)** Prediction of base pair probability by dot and circus plots. The dot plot matrices illustrating the base-pair probabilities with darker dots indicating more chance of base-pairing probability. The circos plot depicting the probability of base -pair for p(WT) and p(MUT) IRX4 RNA sequence, with the sequence initiating from 5’ end at the bottom-left and elongating clockwise reaching to the 3’ end. Each MUT circos plot is demonstrating the variant of interest in red at position 21 at the top. The base pairing of higher probability is highlighted by darker shades of grey. **iii(c1c4)** Another heat map like dot plot matrices representing the base-pairing probabilities between the WT and MUT RNA which is calculated as {Pr(bp in WT) – Pr(bp in mut)}. Strong base-pairing probabilities are denoted by red dots while blue dots are the results of weak base-pairing. The accessibility profile of WT and MUT are denoted in terms of unpaired probabilities. The variation in accessibility (WT-MUT) is indicated by the blue line

### 3.5 Prediction of mutant’s effect on RNA features

The impact of variations on the different features of RNA were represented by base pairing probabilities dot plot, circos plot base pairing probabilities, differential base pairing probabilities dot plot and accessibility profile. The heat map like dot plot matrices with darker dots in MUTs illustrating higher base pairing probabilities (Figure 2ii(b1-b4). Likewise, the circus plot indicating the interconnection between different positions of RNA sequence and their interactions. The arcs that represents the variants connecting the affected nucleotides and showing the potential disruption caused by both the variants (p.Ser24Asn and p.Thr217Iso) using different hues of grey, with darker grey color showing potentially stronger base pairing probabilities (Figure 2iii(c1-c4). The differential dot plot matrices are the difference in base pairing patterns between WT and MUT-RNAs at specific positions. Remarkable changes in the differential dot plot were noted in response to both the missense (p.Ser24Asn and p.Thr217Iso) variants indicating significant deviation in RNA structures due to mutations. Further, the RNA accessibility profile (the probability of being unpaired at each base position which affect the RNA-RNA or RNA-protein interactions) which is crucial for translation was also checked. Both the variants (p.Ser24Asn and p.Thr217Iso) induce notable changes in RNA accessibility.

### 3.6 Analysis of physicochemical properties

The individual amino acid of proteins possesses different properties and any change in the sequence of amino acid of any protein greatly affect its physicochemical properties and thereby also responsible for different secondary and tertiary structures of proteins. The R-side chain of amino-acids greatly define the different physicochemical properties such as alpha-helix, relative mutability, recognition factors and total beta strands of proteins. The missense mutations change the amino-acid, the R-side chain of the newly introduced amino acid will be different from that of the replaced one. We had employed computational approach by Protscale for comparative analysis of these properties for IRX4 proteins. The prediction analysis for recognition factor clearly depicted significant changes induced by both the variants (p.Ser24Asn and p.Thr217Iso). Likewise, comparable changes in relative mutability was noted in response to variant p.Ser24Asn, however negligible alteration caused due to variant p.Thr217Iso. Further, remarkable alterations were observed in total beta strand of both the MUT proteins (p.Ser24Asn and p.Thr217Iso) Figure 3(i).

**Figure 3.**
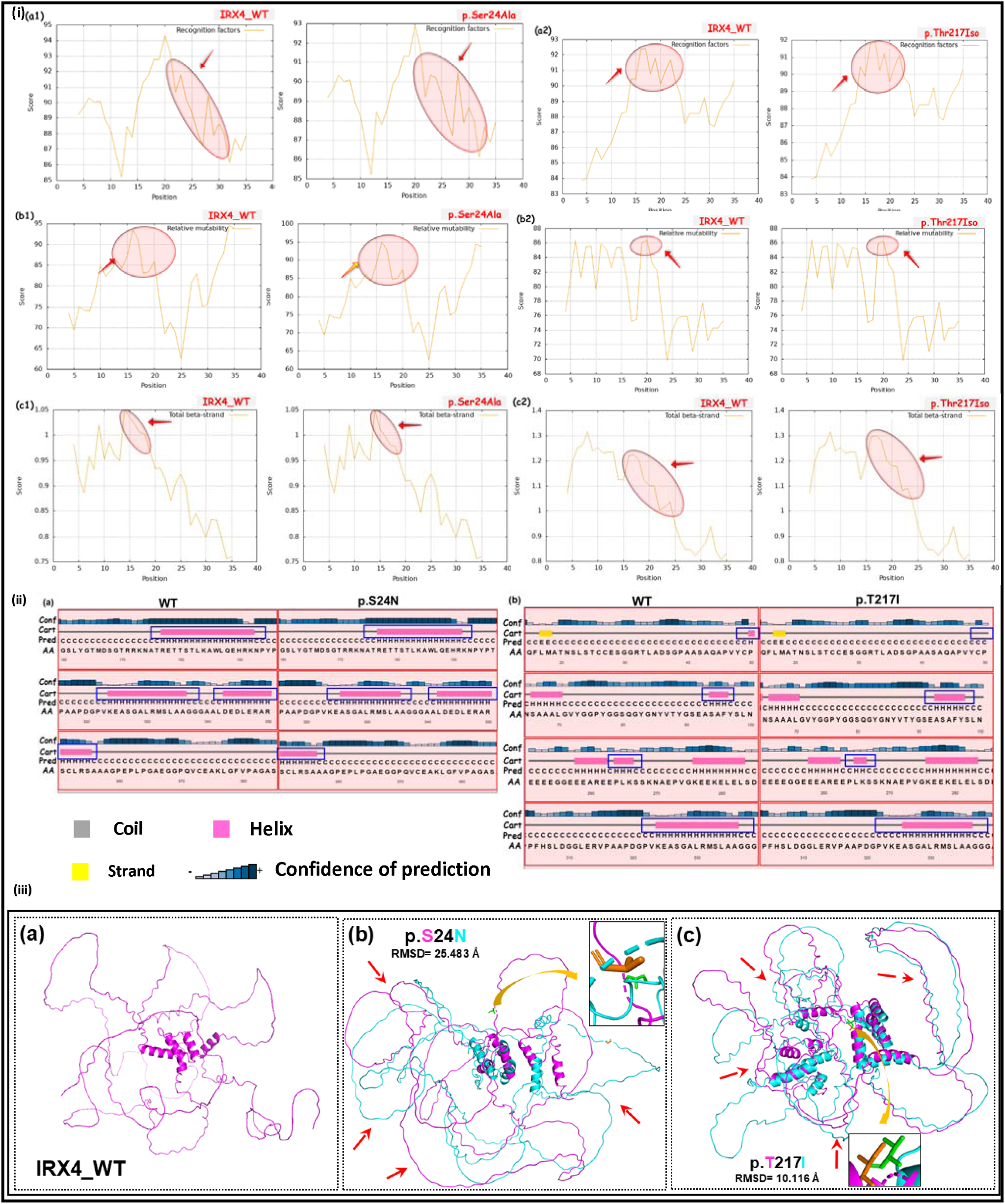
**(i)** Physio-chemical property analysis of IRX4 WT and muteins by computational methods. (a1-a2) comparative analysis of recognition factors of WT and both the muteins, (b1-b2) showing tendencies of relative mutability and, (c1-c2) prediction of total -beta strand. The encircled regions marked with arrow representing the changes. WT= wild-type. **(ii-a-b)** Secondary structures of IRX4 WT and muteins (p.Ser24Asn and p.Thre217Iso) as predicted by Psipred server. The alterations in structures-helix (pink boxes) and strand (yellow boxes) has been highlighted with blue rectangle in both WT and MUTs. (iii) Tertiary structural comparison of IRX4 muteins with WT, (iii-a) modelled-tertiary structure of IRX4 WT, **(b-c)** superimposed tertiary structures of both the muteins of IRX4 (p.Ser24Asn and p.Thre217Iso) over WT with supported global alpha carbon RMSD values. The tertiary structure of WT is shown in magenta colour while muteins are represented in cyan blue colour with mutated residues in sphere model

### 3.7 Predicted alterations in the secondary and tertiary structures of IRX4 muteins

The secondary structures analysis demonstrated significant modifications in the α-helix of muteins. Shortening of α-helix was depicted from position 177-191aa and 316-335aa and extension from aa position 343-350 and 351-360 in mutein p.Ser24Asn. Moreover, α-helix was lost at position 50aa along with extension from aa position 94-97 and shortening from position 265-266aa and 326-336aa. These α-helix changes in secondary structures of muteins were compared to WT (Figure 3ii(a-b).

The superimposed tertiary structures of IRX4-WT and both the muteins depicted structural changes caused by both the variants (p.Ser24Asn and p.Thr217Iso). Significant change was noted in case of mutein p.Ser24Asn with supported RMSD value of 25.483 Å. Similarly, the substitution of threonine by isoleucine also illustrated significant change which was further supported by RMSD value of 10.116 Å (Figure 3iii(a-c).

### 3.8 Effect of variants on the expression and localization of IRX4 proteins

Immunocytochemistry was performed to check the effect of variants on IRX4-WT and IRX4-MUT proteins in H9c2 cell lines. *IRX4* is a transcription factor and expected to localize in nucleus. The result indicated that both the WT and MUT proteins were localized in nucleus and no significant change in the expression of muteins was observed (Figure 4(a1-d3).

**Figure 4.**
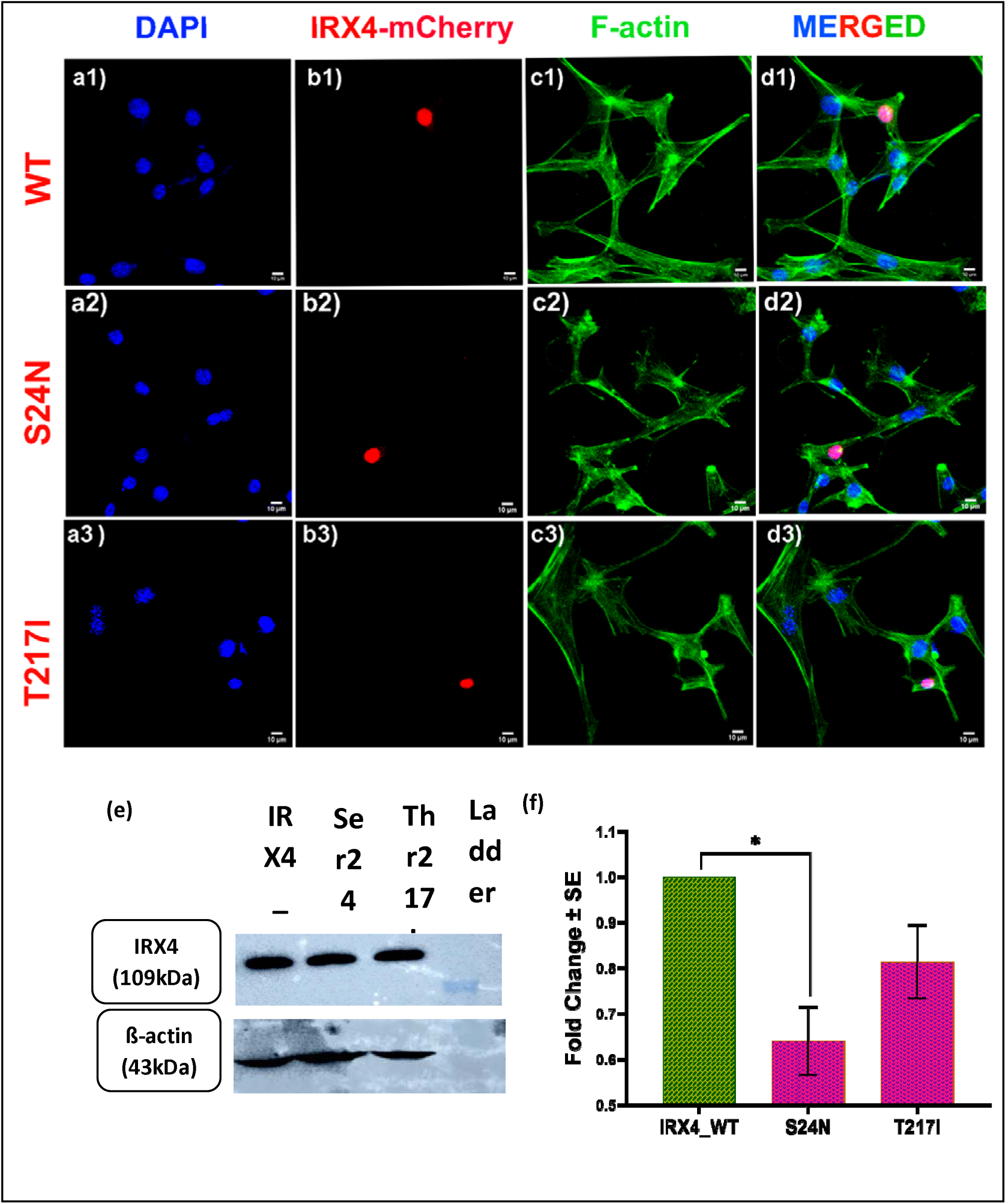
Expression analysis of IRX4 wild-type (WT) and muteins (MUT) by immunostaining and Western blotting. **(a1-d3)** Immunostaining showing cellular location of IRX4 WT and MUT (p.Ser24Asn and p.Thre217Iso) in H9c2 cells. **(e & f)** Western blotting illustrating the expression of IRX4 WT and both the variants (p.Ser24Asn and p.Thr217Iso). *Denotes statistical significance as *\*p<0.05; **p<0.01; ***p<0.001; ****p<0.0001* by one-way ANOVA followed by Dunnett’s multiple comparisons test, *n=3*

### 3.9 Impact of variants on the expression of IRX4 protein estimated by immunoblotting

The expression level of IRX4 protein was checked in response to both the variants (p.Ser24Asn and p.Thr217Iso) in P19 cells. A significant decrease in the expression of IRX4 was reported due to p.Ser24Asn (1.563 fold, p=0.2064) variant while p.Thr217Iso variant revealed decrease in the expression of IRX4 protein albeit not significant (1.229 fold, p=0.0460) (Figure 4(e-f).

### 3.10 Effect of variants on transactivation of Nanog and HEY2 luciferase reporter

To investigate the functional differences of IRX4-MUT proteins vs the IRX4-WT, *in vitro* dual reporter assay was performed with cardiac specific downstream promoters of *IRX4* i.e. *Nanog-luc and HEY2-luc.* The transcriptional activity of each promoter (*Nanog-luc and HEY2-luc*) were normalized with the basal activity of these promoters. The impact of *IRX4*-MUTs on the transcriptional activity of these promoters was calculated as fold change by comparing with the WT. IRX4-WT protein significantly transactivate both the promoters *Nanog-luc and HEY2-luc* by 10.49 fold (p<0.0001) (Figure 5i(a) and 6.403 fold (p=0.0003) respectively (Figure 5i(b). A significant decrease in the transactivation of *Nanog-luc* promoter was observed in response to both the variants p.Ser24Asn by 3.511 fold (p=0.0002) and p.Thr217Iso 3.420 fold (p=0.0002) (Figure 5i(a). Moreover, notable decrease in the activity of *HEY2-luc* was observed due to variant p.Ser24Asn (1.455 fold, p=0.0705) while the TALE-HD domain variant significantly decrease the activity of *HEY2-luc* (2.200 fold; p=0.0044) (Figure 5i(b).

**Figure 5.**
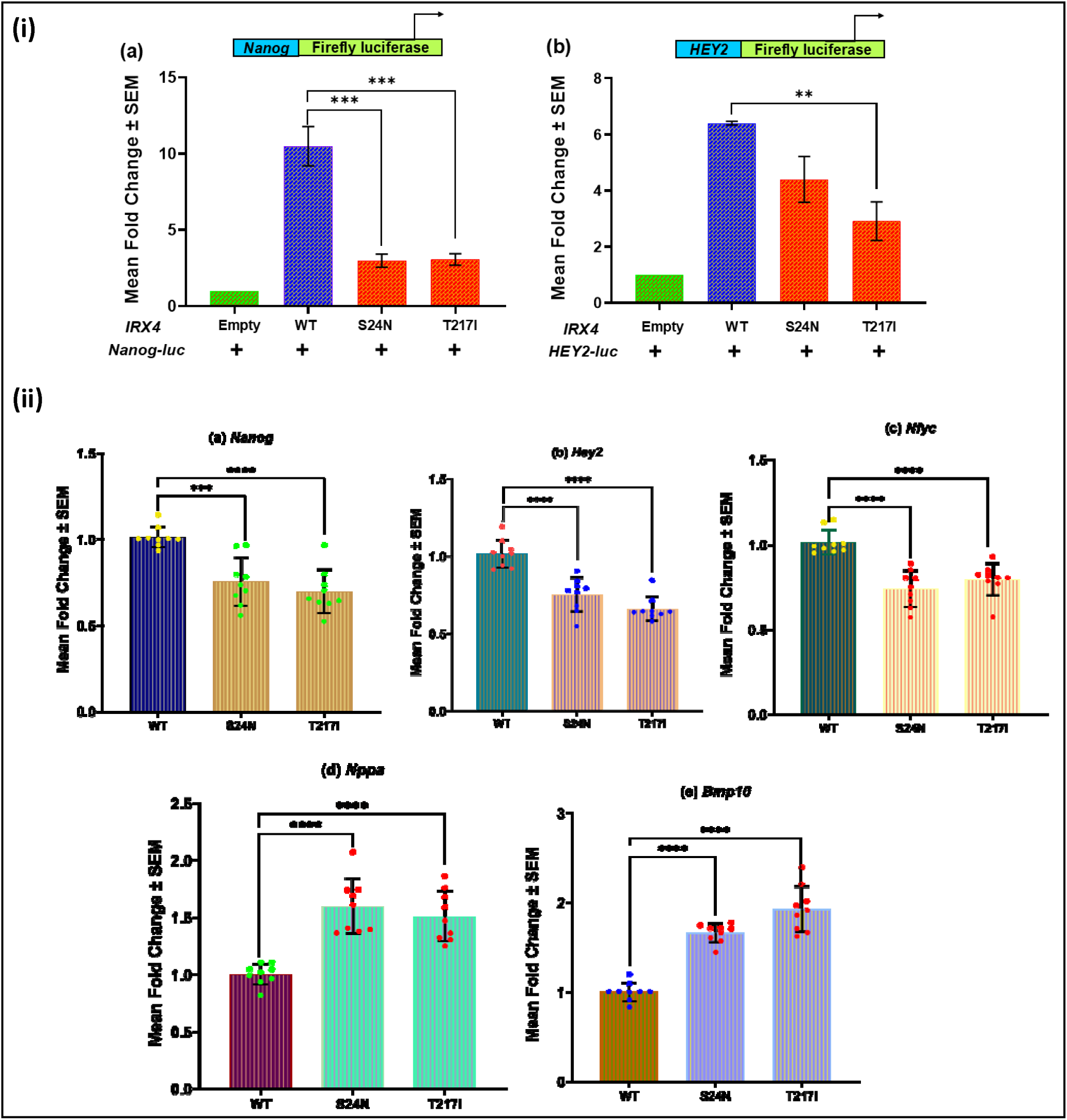
**(i)** Measurement of transcriptional activity of *Nanog-luc* and *HEY2-luc* by dual luciferase reporter assay **(a)** Transcriptional activity of *Nanog-luc* in response to IRX4 WT and MUTs (p.Ser24Asn and p.Thre217Iso) **(b)** Transactivation of *HEY2-luc* due to IRX4 WT and MUTs (p.Ser24Asn and p.Thr217Iso). **(ii)** Graphs illustrating the effect of IRX4 variants (p.Ser24Asn and p.Thre217Iso) on the expression of different downstream targets assessed by RT-PCR **(a)** *Nanog*, **(b)** *Hey2*, **(c)** *Nfyc*, **(d)** *Nppa* and **(e)** *Bmp10.* *Denotes statistical significance as *p<0.05; **p<0.01; ***p<0.001; ****p<0.0001 by one-way ANOVA followed by Dunnett’s multiple comparisons test, *n=9*

### 3.11 Effect of IRX4 variants on the expression of downstream target of IRX4

By real-time PCR, the impact of variants on the endogenous expression of different cardiac specific downstream targets of *IRX4* was examined. The expression of *Nanog* was significantly decreased in response to both the variants p.Ser24Asn (1.325 fold, p=0.0002) and p.Thr217Iso (1.432 fold, p<0.0001) (Figure 5ii(a). Moreover, a significant downregulation in the expression of *Hey2* was induced due to p.Ser24Asn (1.33 fold, p<0.0001) and p.Thr217Iso (1.512 fold, p<0.0001) variants (Figure 5ii(b). The reduced expression of *Nanog* and *Hey2* can be correlated with the luciferase assay data where the expression of both the genes was reduced due to both the variants. Interestingly, significant downregulation of *Nfyc* was noted in response to p.Ser24Asn by 1.349 fold (p<0.0001) and p.Thr217Iso by 1.256 fold (p<0.0001) (Figure 5ii(c). Conversely, there was significant increment in the expression of *Nppa* by 1.6 fold (p<0.0001) and 1.5 fold (p<0.0001) in response to p.Ser24Asn and p.Thr217Iso variants respectively (Figure 5ii(d). Likewise, the expression of *Bmp10* was increased due to both the variants of *IRX4* (p.Ser24Asn by 1.661 fold, p<0.0001 and p.Thr217Iso by 1.929 fold, p<0.0001) (Figure 5ii(e).

## 4. Discussion

*IRX4,* a gene encoding ‘Iroquois-class homeodomain protein,’ is located on chromosome 5p15.33 (HGNC:6129), which produces approximately 2.4 kb transcript that include 7 exons (6 protein coding exons) and 6 introns. The encoded protein is 545 amino acid (aa) long belongs to the family of TALE homeobox transcription factor with TALE-HD (187-225aa), putative transactivation (TAD) (250-268aa) and IRO (361-378aa) domain.

In the present study, the coding region and splice sites of *IRX4* gene were screened in 205 non-syndromic CHD probands, 24 DS with CHD and 27 DS without CHD. Two disease-causing missense variants (p.Ser24Asn and p.Thr217Iso) were identified exclusively in non-syndromic CHDs. Additionally, one known SNP (p.119Ala>Thr) was identified in non-syndromic CHD as well as CHD with DS. Both the missense (p.Ser24Asn and p.Thr217Iso) variants were not reported in any variant database and were not detected in the screening of 300 healthy chromosomes. The first novel variant (p.Ser24Asn) and the reported SNP (p.119Ala>Thr) confined to the N-terminal region (encoding amino acids 1-135) of *IRX4*. This region is crucial for the interaction between *Irx4* and *Rxra* to regulate the expression of ventricular genes (Wang et al. 2001). The other novel variant (p.Thr217Iso) lie in the TALE-HD which facilitate in DNA binding. Although analysis on their functional characterization are scanty, only one study reported N-terminal (p.Asn85Tyr and p.Glu92Gly) variants caused functional deficit in IRX4 protein. Interestingly, the functional impact of TALE-HD variants has not yet been explored. Hence, this study performed *in silico* and *in vitro* functional analyses of these novel variants to elucidate their potential association with the disease.

The amino acid substitutions in both variants (p.Ser24Asn and p.Thr217Iso) are highly conserved among mammalian species and were absent in the screening of 300 healthy chromosomes, indicating their potential pathogenicity. Moreover, the majority of *in silico* prediction tools classified these variants as damaging or disease-causing, which could be correlated with the pathogenesis of CHD.

Moreover, the mRNA structural modifications caused by the p.Ser24Asn and p.Thr217Iso variants are believed to influence RNA folding. This is supported by analyses of base pairing probabilities and accessibility profiles, which suggest alterations in RNA stability and functionality. Such changes may interfere with normal molecular processes, potentially leading to disease (Salari et al. 2013).

As a consequence, the p.Ser24Asn and p.Thr217Iso variants showed notable differences in several physicochemical properties such as recognition factors, relative mutability, and overall beta strand content which are thought to influence protein folding, stability, structure, and molecular interactions (Boon et al. 2020). Interestingly, predictions of their secondary structures indicated changes in the alpha-helix, suggesting a disruption in protein architecture and its ability to interact with other molecules. The notable modifications in β-sheet disordered the tertiary structures which was also depicted by the tertiary structural change analysis.

Furthermore, our protein assay using Western blotting showed decreased expression levels for both the N-terminal and TALE-HD variants (p.Ser24Asn and p.Thr217Iso). This reduction is likely due to altered RNA secondary structure or the production of unstable and misfolded proteins, which may impair the ability of these muteins to interact effectively with other molecules.

A study by Jia and group (2020) reported that *IRX4* knockdown significantly impaired *NANOG* transcriptional activity (Jia et al. 2020; Waisman et al. 2024). Similarly, reporter gene assays in the present study revealed that both the N-terminal and TALE-HD muteins (p.Ser24Asn and p.Thr217Iso) caused a significant decrease in *Nanog* promoter activity. *Nanog* is a known cellular marker, expressed in myocardial cells, fibroblasts, and small round cells across different myocardial regions (Luo et al. 2014). The reduced *Nanog* expression observed with both muteins may be linked to disruptions in upstream signaling pathways. Furthermore, the enhancer activity of *HEY2* was also markedly reduced by the same muteins. *Irx4* has been shown to work synergistically with *Hey2* to suppress atrial gene expression in the ventricular myocardium (Xin et al. 2007; Wang et al. 2024) and mutations in *IRX4* are thought to impair its interaction with HEY2 (Cheng, 2011). *Hey2* is specifically expressed in ventricular myocardium and plays a critical role in regulating ventricular gene expression (Leimeister et al., 1999; Xin et al., 2007). We hypothesize that structural changes in the muteins may interfere with their cooperative binding to HEY2, potentially disrupting downstream signaling pathways. Notably, previous findings also support our observations, showing that N-terminal variants (p.Asn85Tyr and p.Glu92Gly) reduced promoter activity by impairing DNA-binding affinity (Cheng, 2011). However, to further investigate the functional consequences and assess DNA-binding and protein-protein interactions of these variants, pull-down assays or co-immunoprecipitation (co-IP) experiments are recommended.

Similarly, qRT-PCR gene expression analysis revealed a significant decrease in the expression of *Nanog* which is a downstream effector of *IRX4* due to both the N-terminal and TALE-HD variants (p.Ser24Asn and p.Thr217Iso). Likewise, a significant reduction in the expression of *Hey2* which is regulated by *IRX4* in the ventricular myocardium (Xin et al. 2007; Wang et al. 2024) in response to both the N-terminal and TALE-HD variants (p.Ser24Asn and p.Thr217Iso) was observed. This supports our previous reporter gene analysis findings. Besides this, the expression of another transcription factor crucial for cardiogenesis *Nfyc* (Li et al. 2015; Dolfini et al. 2024) indicated a significant decrement in the expression caused by both the variants (p.Ser24Asn and p.Thr217Iso). Previous studies have shown that *IRX4* knockdown results in reduced *NFYC* activity (Vitharanage 2023). In contrast, cardiac specific transcription factor *Nppa* was significantly overexpressed in response to both the variants (p.Ser24Asn and p.Thr217Iso). The enhanced expression of *Nppa* is correlated with study of Bruneau et al. 2001 who investigated that lack of *Irx4* mice embryo results in the overexpression of *Nppa* in the ventricles (Bruneau et al. 2001). Correspondingly, increment in the expression of *Bmp10* has also been underscored due to both the variants (p.Ser24Asn and p.Thr217Iso). Overexpression of *Bmp10* is supported by the elevated level of *Bmp10* in *Irx3*;*Irx4* double knockout hearts (Liu 2017) and direct repression of *Bmp10* in endocardium by *Irx* genes (Wang et al. 2024).

We observed a heterogenous phenotype of CHD namely VSD; DORV, TA, ASD; Cyanotic CHD in N-terminal variant (p.Ser24Asn). This suggests that *IRX4* plays a critical role at various stages of heart development, beyond just interventricular septum formation. A collaborative study by Vincent M. Christoffels (2000) in mice explored the dynamic expression of *Irx4* at different developmental stages, further supporting this observation (Christoffels et al. 2000). In contrast, the HD variant (p.Thr217Iso) was specifically linked to VSD, implying that this variant might uniquely impact *IRX4* involvement in ventricular septum formation. Our *in-silico* analysis, examining changes in structural and physicochemical properties, strongly suggests that both variants are likely disease-causing, potentially disrupting *IRX4* interaction with other molecules. Moreover, *in vitro* functional characterization, including Western blotting, revealed reduced expression of both muteins. Gene expression analyses of *IRX4* interacting partners (*Hey2* and *Bmp10*) and downstream effectors (*Nppa*, *Nanog*, and *Nfyc*) further supported these findings. Transactivation assays with *Nanog* and *HEY2* provided additional evidence that IRX4 binding is impaired, disrupting downstream signaling pathways due to both variants. Overall, we could infer that our functional analysis strongly demonstrate that both the variants are pathogenic and have adverse effect on the developmental pathway of septation. Besides, it also interfering with other crosstalk signalings which are crucial for multiple stages of cardiogenesis.

It is noteworthy that the VSD phenotype has been previously reported in the *IRX4* gene in the Chinese population, with functional implications suggesting its potential causative role in the development of the interventricular septum (Cheng et al. 2011). However, mice lacking *Irx4* did not display the VSD phenotype, which may be attributed to species differences in the function of *IRX4*. Alternatively, other unidentified factors could be interacting with *IRX4*, modifying its phenotypic expression.

No disease-causing variants in *IRX4* were identified in CHD with DS, suggesting that this gene may not be directly associated with CHD in individuals with DS. However, a reported SNP was found specifically in CHD with DS (not in DS without CHD). Previous studies have indicated that overexpression of VEGF pathway genes due to trisomy 21 disrupts epithelial– mesenchymal transformation, leading to septal defects (Ackerman et al. 2012; Marder et al. 2015). Additionally, other studies have highlighted the potential involvement of Hedgehog (Ripoll et al. 2012), calcineurin (Fuentes et al. 2000), and folate pathways (Locke et al. 2010) in the pathogenicity of CHD with DS. While trisomy 21 is recognized as a risk factor for CHD, it is important to note that only 40-60% of individuals with DS are affected by CHD (Pradat 1992; Stoll et al. 1998b; Freeman et al. 1998).

## 5. Conclusion

The findings of our study infer that *IRX4* variants are linked with diverse phenotype of CHD in non-syndromic cases of CHD only. The N-terminal variant is associated with mild to severe phenotype which underpin the involvement of *IRX4* at various stages of cardiogenesis along with inclusion in ventricular septum formation. Our functional characterization of variants also strengthens in unveiling the crucial role of N-terminal region in recognizing the DNA binding sites. Besides, the TALE-HD variant also delineating the relevance of the domain. The experimental studies somewhat provide novel insights in unravelling the molecular mechanism of the disease which implying the disease-causing potential of *IRX4*. However, to further elaborate the role of *IRX4* with domain-wise function, more functional studies and screening need to be performed. Our findings revealed no pathogenic *IRX4* variants in CHD with DS, indicating that the gene may function via a mechanism independent of trisomy 21-associated pathways.

## Acknowledgements

We are grateful to all the patients, their family members and control individuals for their participation in the present study. We would like to acknowledge Dr. Ramkumar Sambasivan (InStem, Bengaluru, Karnataka, India) for providing P19 cells, Dr. Yusuke Watanabe from National Cerebral and Cardiovascular Center Research Institute, Suita, Osaka, Japan for *HEY2* luciferase reporter construct and Dr. Ariel Waisman from Institute of Neuroscience (INEU), CONICET, Buenos Aires, Argentina for *Nanog* luciferase reporter construct. We would like to extend thanks to University Grants Commission (UGC) for research fellowship (JRF and SRF) to Jyoti Maddhesiya.

## Credit author statement

**Bhagyalaxmi Mohapatra:** Conceptualization, supervision, investigation, formal analysis, data curation, writing original draft and editing, funding acquisition, and project administration.

**Jyoti Maddhesiya:** Investigation, methodology, data curation, validation, software, formal analysis, writing original draft, writing-review and editing.

**Kriti Jain:** Methodology

**Dharmendra Jain:** Clinical investigation and patient enrolment

**Ashok Kumar:** Clinical investigation and patient enrolment

**Amit Rai:** Clinical investigation and patient enrolment

## List of abbreviations

CHD: congenital heart disease
DS: Down syndrome
AVSD: atrioventricular septal defect
ASD: atrial septal defects
VSD: ventricular septal defects
DORV: double outlet right ventricle
TA: tricuspid atresia
BAV: bicuspid aortic valve
PDA: patent ductus arteriosus
LVNC: left ventricular non-compaction

