## Supplementary Information for "Unravelling the role of *IRX4* variants in non-syndromic and Down syndrome associated congenital heart disease"

**Full blot images of Western Blots**

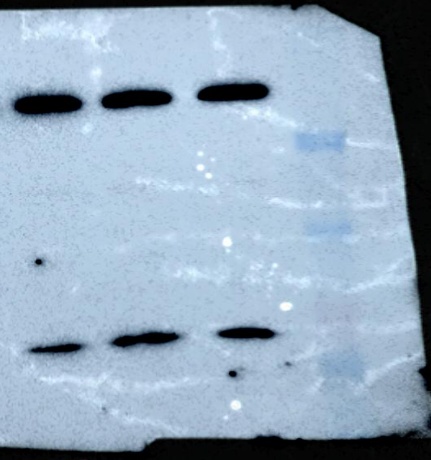

**IRX4 (109kDa)**

**IRX4_WT**

**S24N**

**T217I**

**Beta-actin (43kDa)**

**Ladder**

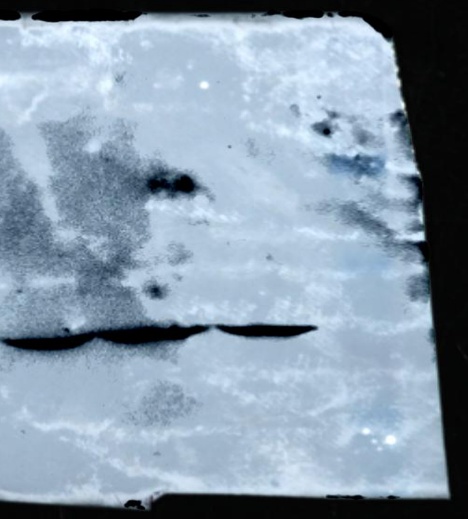

***Supplementary Fig. s1.*** Full-length, uncropped Western blot images showing IRX4, and β-actin protein expression. The images presented in the manuscript are the cropped images of these blots.

**Supplementary Table s1** *Clinical phenotype with their relative frequency in male and females*

| **CHD** | **CHD Phenotype** | **Number (%)** |
| --- | --- | --- |
| **Acyanotic defects** | Ventricular septal defects (VSD) | 65 (31.7) |
|  | Atrial septal defects (ASD) | 36 (17.56) |
|  | ASD, VSD | 16 (7.8) |
|  | Patent ductus arteriosus | 18 (8.7) |
|  | Patent foramen ovale | 5 (2.43) |
| **Cyanotic defects** | Tetralogy of Fallot | 25 (12.19) |
|  | Transposition of great arteries | 5 (2.43) |
|  | Dextrocardia | 2 (0.97) |
|  | Tricuspid Atresia | 4 (1.95) |
|  | Double outlet right ventricle | 3 (1.46) |
|  | Pulmonary Atresia | 3 (1.46) |
|  | Total Pulmonary Venous Connection | 1 (0.48) |
| **Left/ right obstructive defects** | Bicuspid aortic valve | 2 (0.97) |
|  | Pulmonary stenosis | 18 (8.7) |
|  | Aortic stenosis | 1 (0.48) |
|  | Coarctation of aorta | 1 (0.48) |
| **Total** | | **205** |

**Supplementary Table s2**. *qRT-PCR primer Sequence with amplicon length and annealing tempearture*

| **Gene** | **5’ Sequence 3’** | **(Ta) (°C)** | **Amplicon size (bp)** |
| --- | --- | --- | --- |
| *Nanog* | FP- AAGCAGAAGTACCTCAGCC | 60 | 228 |
|  | RP- CAGATGCGTTCACCAGATA |  |  |
| *Hey2* | FP- CAGTGATGAGGTCCAATTC | 54 | 136 |
|  | RP- GCTGTTGGCACTAGTCTTC |  |  |
| *Nfyc* | FP- GACCACCAGTTCTACGACC | 56 | 204 |
|  | RP- GTGTTGGTAATGATCTGCT |  |  |
| *Nppa* | FP- GCTTCCAGGCCATATTGGAG | 54 | 126 |
|  | RP- GGGGGCATGACCTCATCTT |  |  |
| *Bmp10* | FP- GTTTCTCAAGACGCTGAAC | 58 | 205 |
|  | RP- GAGGAGAGGATATTTCCGGA |  |  |

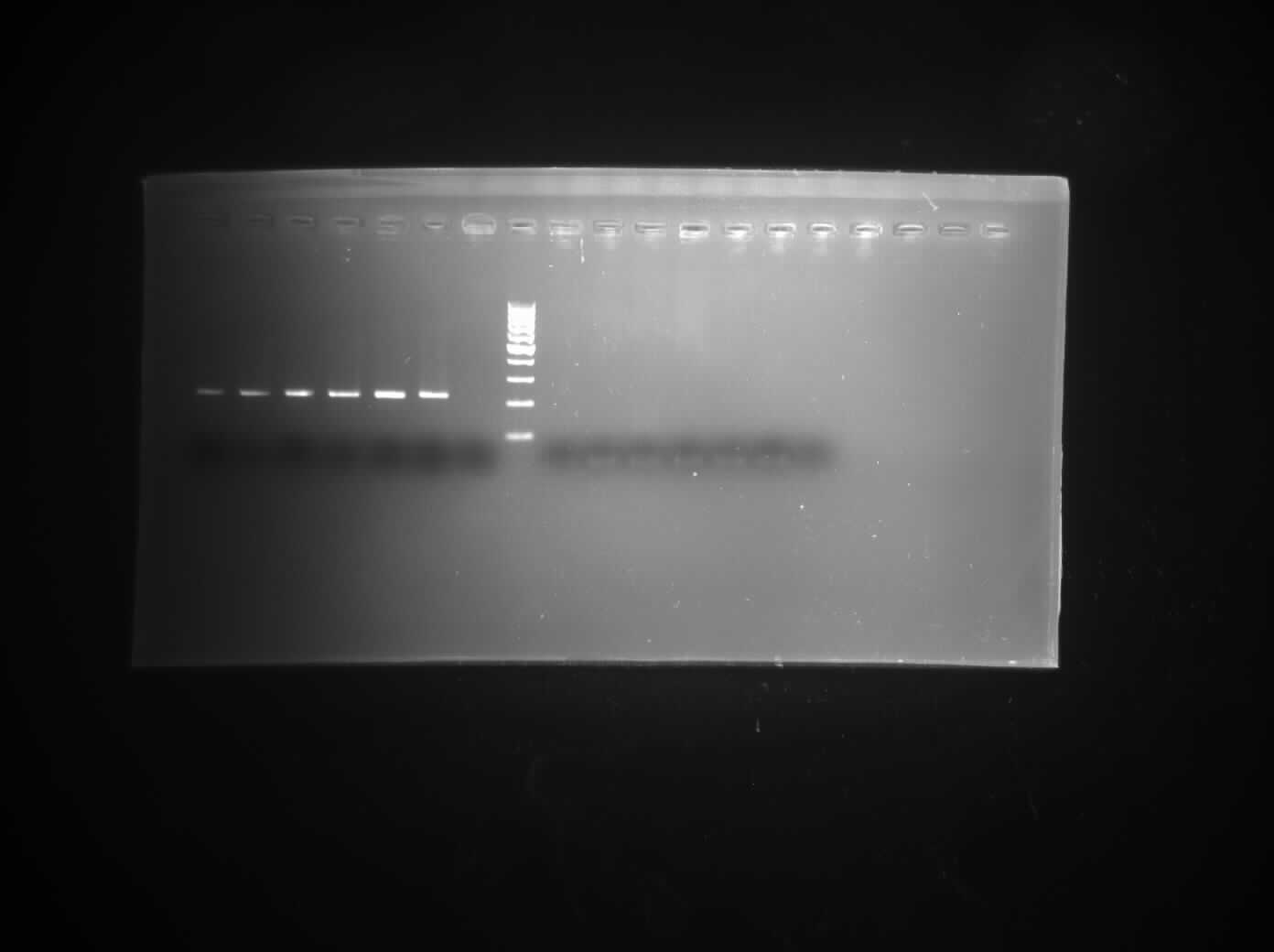

**52	54	56	58	60	62	-ve	L**

***Nanog***

**(228bp)**

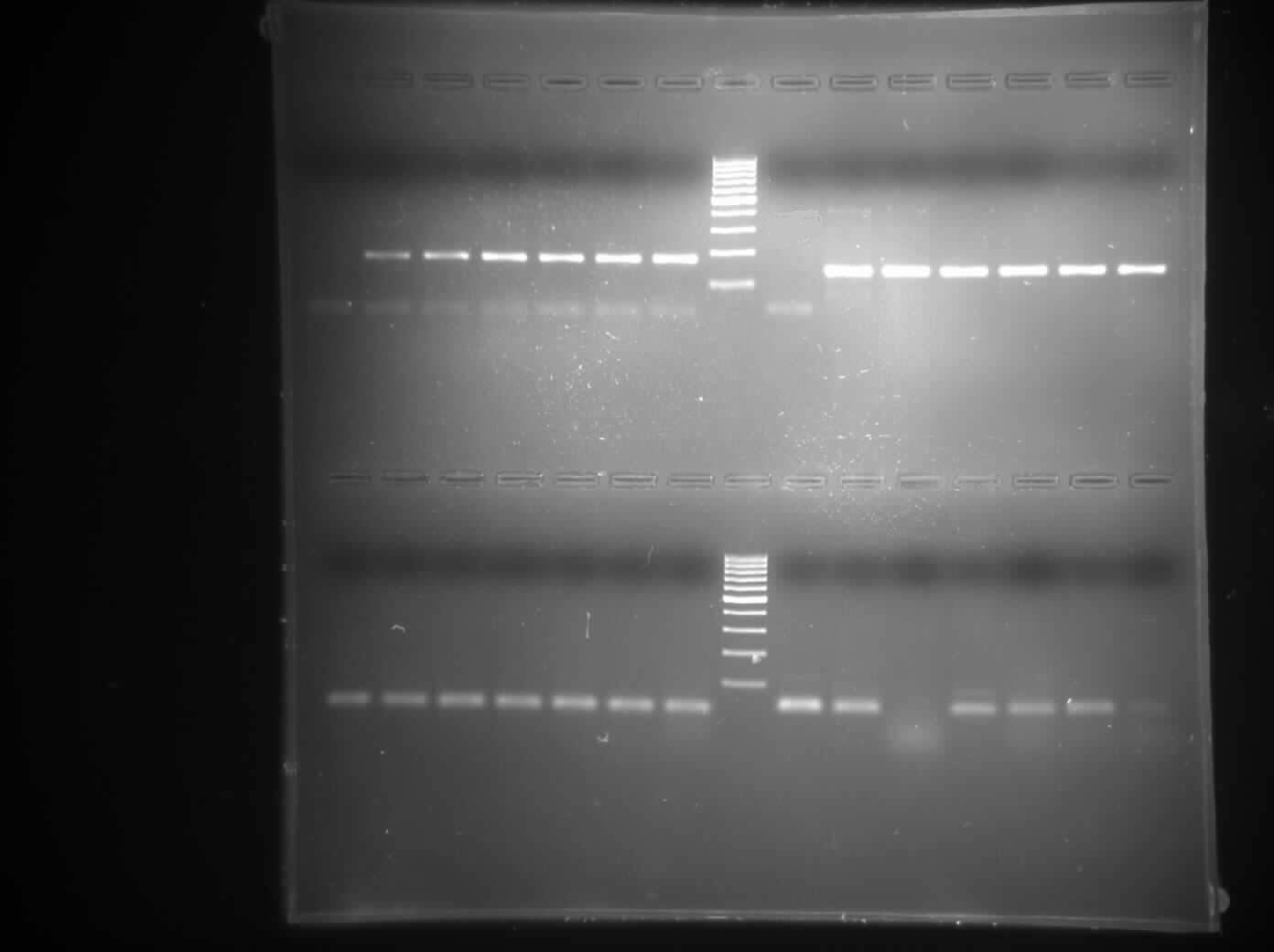

**L	-ve	52	54	56	58	60	62**

***Hey2***

**(136bp)**

***Nppa***

**(126bp )**

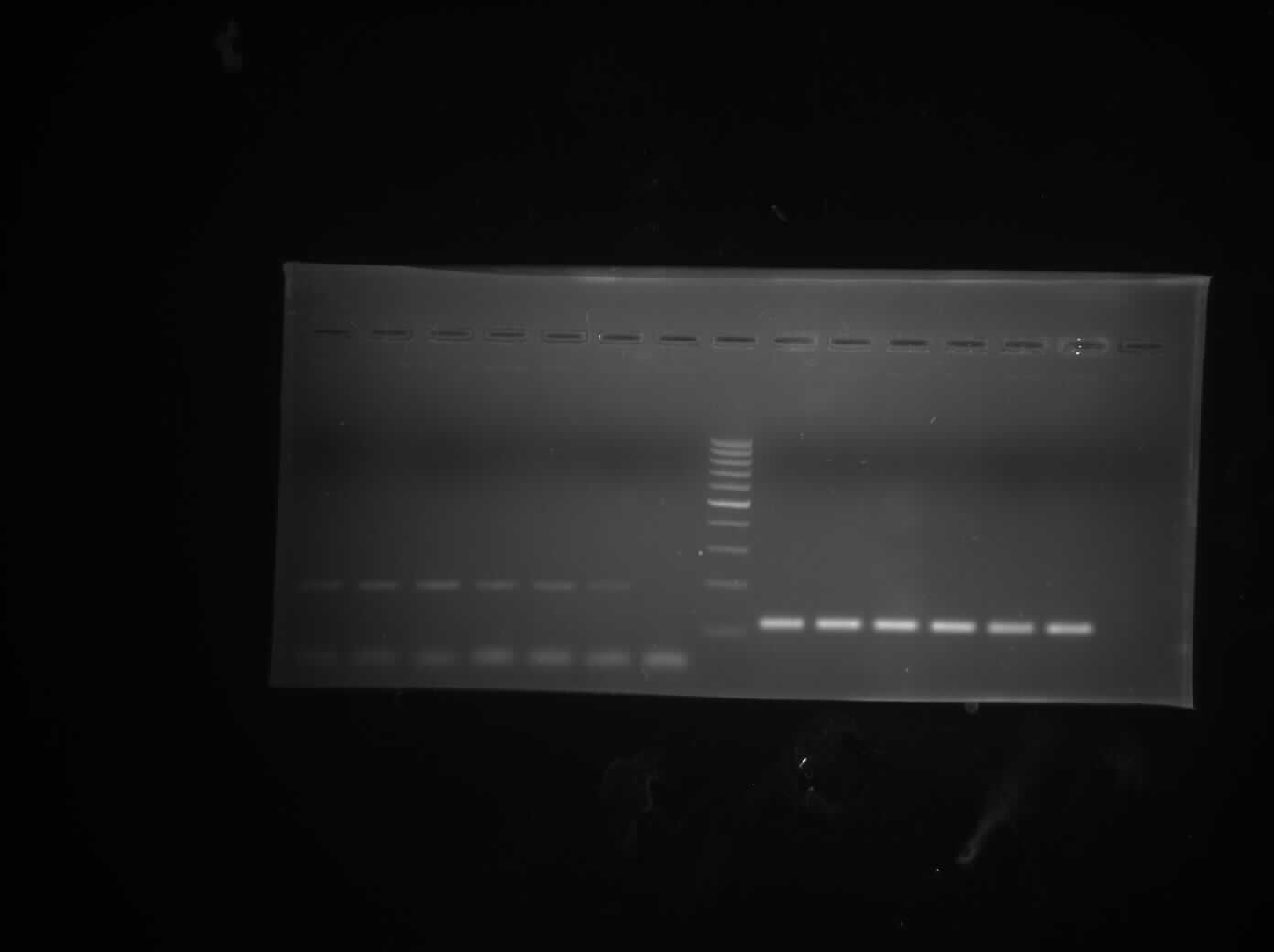

**L	52	54	56	58	60	62	-ve**

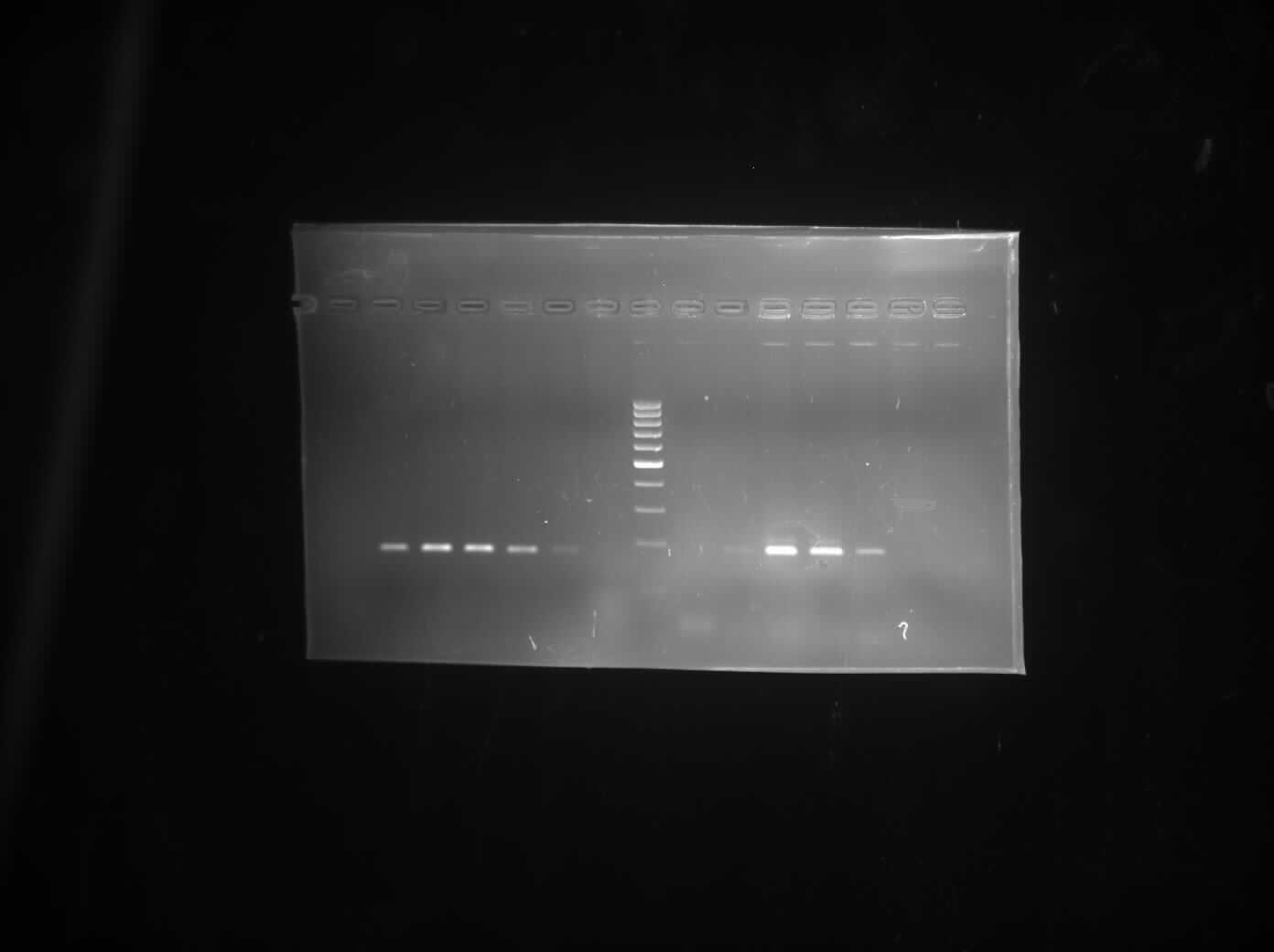

**L	-ve	52	54	56	58	60	62**

***Nfyc***

**(204bp)**

***Bmp10***

**(205 bp)**

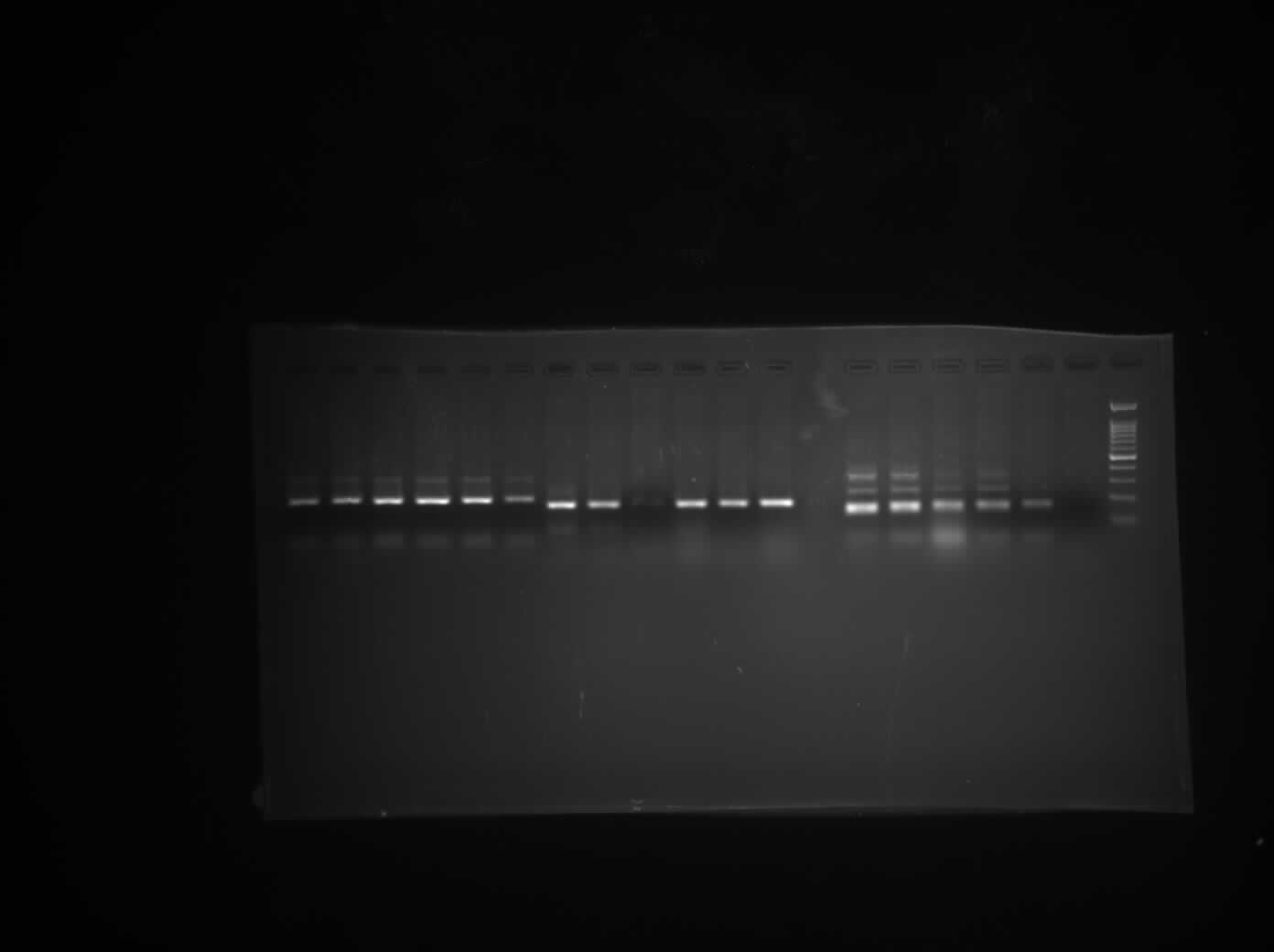

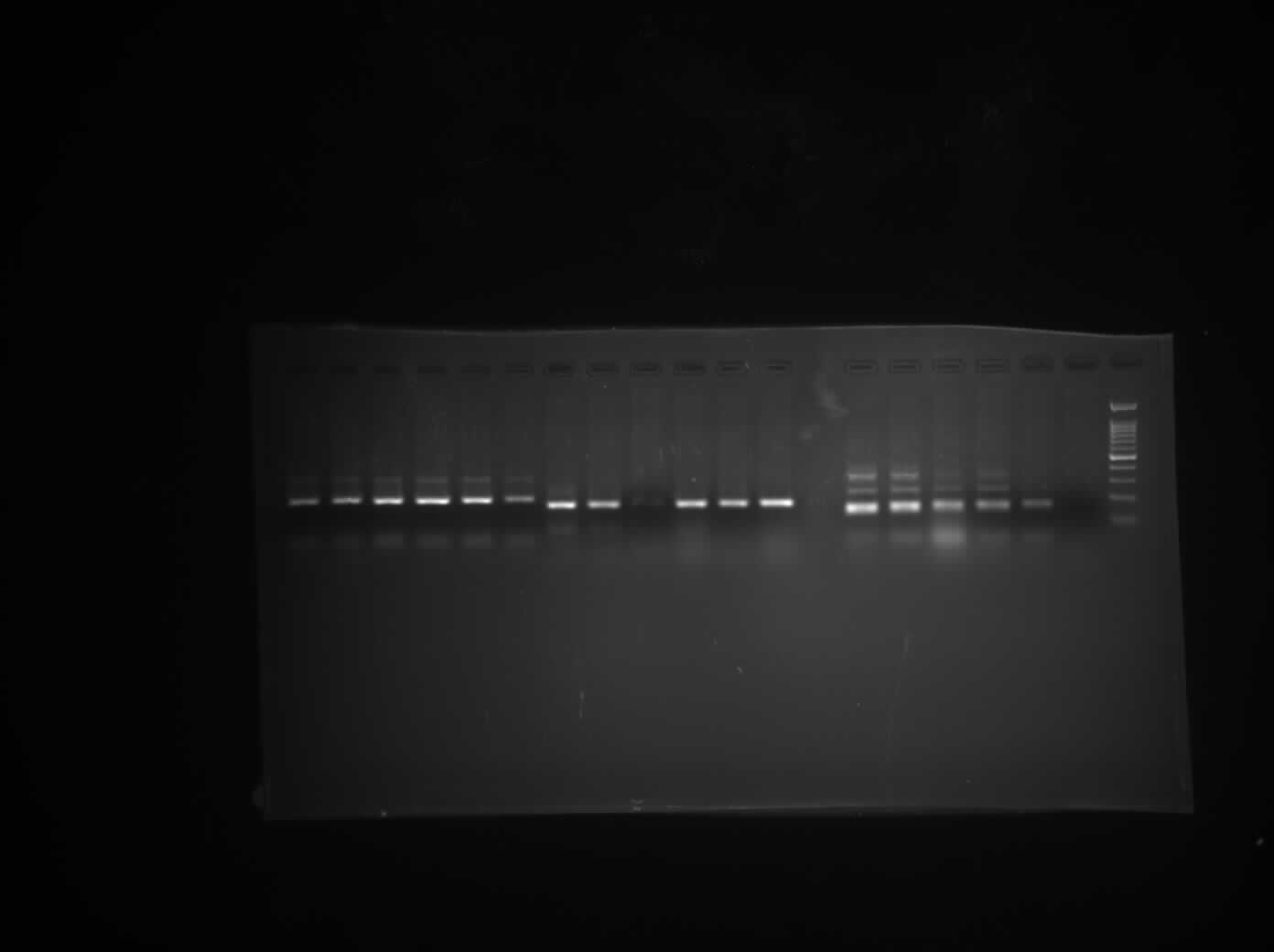

**52	54	56	58	60	62	-ve	L**

***Supplementary Fig s2.*** Semi-quantitative PCR gel images representing the gradient PCR of qRT-PCR primers (*Bmp7, Bmp2, Nkx2.5, Gata4, Mef2c, Irx4, Smad1, Smad4, Smad5, Nodal, Pitx2, Chordin*) standardization from gradient temperature 52-62**°**C. The amplicon length was determined against a **100 bp DNA ladder** as the molecular size mark
